# Mycotoxin-driven proteome remodeling reveals limited activation of *Triticum aestivum* responses to emerging chemotypes integrated with fungal modulation of ergosterols

**DOI:** 10.1101/2025.06.30.662327

**Authors:** Seyedehsanaz Ramezanpour, Nasim Alijanimamaghani, Jason A. McAlister, David Hooker, Jennifer Geddes-McAlister

## Abstract

Fusarium head blight (FHB), mainly caused by *Fusarium graminearum*, is a globally important wheat disease reducing yield and grain quality. The pathogen produces mycotoxins deoxynivalenol (DON), 3-acetyl DON (3ADON), 15-acetyl DON (15ADON), and nivalenol (NIV), which threaten food and feed safety. During the past 15 years, surveillance has identified nove trichothecenes 3ANX and NX, which show increased virulence compared to DON. In this study, we investigated the effects of 15ADON/3ANX chemotype on both wheat and *F. graminearum* proteomes to identify proteins and pathways responsive to the emerging mycotoxin chemotype. We defined a core wheat proteome across all strains (15ADON- and 15ADON/3ANX-producing, and untreated controls) to explore changes in protein abundance associated with defense response, grain development, and reduced photosynthesis upon infection. Conversely, we identified 32 wheat proteins exclusively produced in the presence of 15ADON/3ANX strains, providing further insight into chemotype-specific responses of wheat. Additionally, assessment from the fungal perspective, reported 119 proteins exclusive to the 15ADON/3ANX strains, including those associated with virulence and mycotoxin production. Lastly, investigation of strain-specific proteome changes showed a significant reduction in mycotoxin protective mechanisms in wheat upon exposure to two 15ADON/3ANX strains, as well as a novel connection between elevated ergosterol biosynthesis and 15ADON/3ANX producing strains. Together, our study characterizes distinct protein production profiles in wheat and *F. graminearum* in response to 3ANX and provides evidence that these molecular changes influence fungal virulence and host defense responses.

## Introduction

Wheat production is globally important for food production and economies. Within Canada, wheat production exceeds 30 M tonnes annually with 75% of the grain exported to China, Indonesia, Japan, Bangladesh, and the United States, bringing in over $9 B to the economy^1,2^. Given the significant importance of this commodity in the national and economies and the necessity to satisfy the dietary requirements of the expanding population, it is imperative to ensure that wheat products are safe and free from disease-related contaminants. Presently, wheat productivity and marketability are threatened by over 30 pests and pathogens, with Fusarium head blight (FHB) being the most damaging disease affecting wheat yield and quality in the previous three decades ^3–5^. The disease has led to reduced market value due to contamination with harmful mycotoxins, such as deoxynivalenol (DON) ^6–8^. Management strategies to mitigate the impact of FHB in wheat include the use of more resistant cultivars, fungicide applications, and other agronomic practices^9^.

*Fusarium graminearum* is the primary cause of FHB in North America, infecting wheat at flowering through airborne ascospores and splash-dispersed macroconidia, with the fungus surviving on crop residues as mycelium or spores^10^. *F. graminearum* produces numerous types of trichothecene mycotoxins, which are categorized based on the nature of the C-8 substitutes^11,13^. For instance, type-B trichothecenes include DON, 3-acetyl DON (3ADON), 15- acetyl DON (15ADON), and nivalenol (NIV) with a ketone group at the C-8 position, whereas type-A trichothecenes exhibit either a methylene hydroxyl or an ester group at the C-8 position^15,17^. Upon consumption of contamianted food or feed by humans and livestock, respectively, trihothecenes will inhibit protein synthesis in eukaryotic organisms, exhibit cytotoxic properties, provoke vomiting and loss of appetite in livestock, and influence immune system activity^19,20^. In wheat, the production of 3ADON or 15ADON by *F. graminearum* is converted to DON, destroying the membranes of adjacent plant cells, releasing nutrients, and accelerating growth of the fungus ^21–23^.

During the past 15 years, *F. graminearum* strains producing alternative mycotoxins have been collected from the Upper Midwest region of the United States and throughout the Great Lakes area. Although, characteristic symptoms of FHB were described, chemical analysis defined synthesis of a novel trichothecene, 3ANX, and the deacetylated variant, NX, which resembled 3ADON and DON, respectively, lacking the C-8^24–26^. Further surveillance identified an additional population of *F. graminearum*, which could produce both 15ADON and 3ANX^27,28^ The emergence of *F. graminearum* populations that do not produce DON, rather 3ANX, suggests an adaptive response by the fungus to the elimination of 3- or 15-ADON as necessary virulence factors^15^. Moreover, as 3ANX inhibits protein synthesis at similar concentrations to DON and is more toxic than DON within evaluated animal models, investigation into its cytotoxic effects and mechanisms of plant defense modulation are warranted^25,29^.

Given the emergence of new mycotoxin chemotypes produced by *F. graminearum* strains, and the economic importance of wheat in Canada, we aimed to assess protein level differences attributed to the new mycotoxin, 3ANX. Working with strains from Ontario, Canada, we performed a field inoculation trial with 15ADON- and 15ADON/3ANX-producing *F. graminearum* phenotypes over three weeks. We performed mass spectrometry-based proteomics profiling on the wheat heads inoculated with strains from each chemotype and compared wheat proteome changes relative to the chemotype and an uninfected control. We also assessed differences in the fungal proteome relative to each chemotype, and defined strain-specific responses from both the wheat and fungal perspectives. Upon infection, our findings highlighted wheat proteome enrichment of proteins associated with defenses response, chitinase production, and lipid transport, and a reduction in plant photosynthesis. We defined significantly different proteins between chemotype and control; however, no significantly different proteins were observed by comparison between the chemotypes, suggesting that the presence of 15ADON/3ANX does not specifically alter abundance of wheat proteins. Interestingly, we observed exclusive production of wheat proteins specific to each chemotype with an emphasis on protein structure and binding, endopeptidases, and oxidoreductase response in the presence of 3ANX. From the fungal perspective, we observed a 3ANX-exclusive proteome, including proteins associated with fungal virulence and mycotoxin production, highlighting differences in the pathogen correlating with mycotoxin chemotype.

Lastly, we explored strain-specific responses from both the wheat and pathogen perspectives and observed a significant reduction in wheat ß-glucanase production in a single 15ADON strain and a significant reduction in wheat glutathione transferases production in two 15ADON/3ANX strains, suggesting reduced activation of classical plant mycotoxin protective mechanisms in the presence of 3ANX. For *F. graminearum*, we observed reduced production of enolase in two 15ADON strains and elevated production of ergosterol biosynthesis protein (Erg19) in six 15DON/3ANX strains, providing evidence of a novel correlation between ergosterol production and 3ANX. Overall, we define unique protein production profiles for both wheat and fungus in the presence of 3ANX and provide evidence of these molecular profiles influencing fungal virulence and host defense response.

## Materials and Methods

### Fungal culture and macroconidia preparation

*Fusarium graminearum* strains used in this study (Table S1) were cultured on potato dextrose agar (PDA) plates and incubated for 5 d at room temperature in the dark. Preparation of macroconidia suspension was performed as previously described^30^ with some modifications. Bilay’s liquid medium consisted of the following components per liter of distilled water: 8.0 g KNO, 4.0 g KH PO, 2.0 g KCl, 2.0 g MgCl, and 2.0 g sucrose. Media was dispensed at 205 mL per 500 mL Erlenmeyer flask and autoclaved. Ten 1 cm² PDA agar plugs containing a single strain of *F. graminearum* were added to each flask with cultures prepared in triplicate for each strain and incubated on a rotary shaker at 120 rpm and 25 °C for 14 d. One flask containing uninoculated Bilay’s medium was included as a negative control. After three weeks, the spore suspensions were filtered through sterile cheesecloth to remove mycelial fragments and other debris. The macroconidia were verified morphologically and quantified using a hemocytometer. The suspension was adjusted to a concentration of 50,000 macroconidia/mL using sterile distilled water^30,31^.

### Wheat cultivation

The artificially inoculated-misted experiment consisted of 21 strains inoculated in four replicates arranged in a Split Plot Design. Each plot (experiment unit) was 2 m in length by 1.6 m in width, which consisted of 8 rows spaced 18-20 cm (6-7 inches) apart. A total of 172 seeds were planted in each row. The winter wheat cultivar 25R40 (susceptible to FHB), distributed by Pioneer Company (Guelph, ON, Canada) was planted in October 2021 at the Elora Research Station (University of Guelph. Ontario, Canada). Plots were artificially spray-inoculated at 50% anthesis stage using the macroconidia suspension of *F. graminearum* strains^32^. Inoculation was performed in June 2022 followed overhead mist irrigation for 3 weeks after flowering to favor FHB development. Samples were harvested 3 weeks post inoculation with 10 heads per strain pooled into one collection bag. Samples were kept on ice for transport and then stored at −80 °C until protein extraction was performed.

### Protein extraction

Wheat heads were processed as we previously described^33^. Briefly, 10 heads from infected (or uninfected) samples were pooled and ground in liquid nitrogen using a mortar and pestle, followed by resuspension in a lysis buffer (100 mM Tris-HCl, pH 8.5) containing a protease inhibitor cocktail tablet and 2% sodium dodecyl sulfate, and sonicated in a water bath at 4 °C for 30 s on/30 s off for 3 cycles. Then 10 mM dithiothreitol (final concentration) was added and samples incubated at 98 °C for 10 min with 800 rpm rotation. Samples were cooled to room temperature and 5 mM iodoacetamide (final concentration) was added followed by incubation at room temperature in the dark for 20 min. Centrifugation was performed at 9,000 x *g* at 4 °C for 10 min to remove plant debris, and supernatant was transferred to a new tube followed by protein precipitation overnight at −20 °C in 100% acetone. Precipitated proteins were collected at 16,000 x *g* and 4 °C for 10 min, supernatant was removed, and the pellet was washed twice with 80% ice cold acetone. Following air drying of the pellet, samples were resuspended in 8 M urea/40 mM HEPES, sonicated at 4 °C in a water bath at 30 s on/30 s off for 5 cycles. Protein concentration was measured, and samples were diluted with 50 mM ammonium bicarbonate (2 mM final urea concentration) and digested overnight with trypsin/LysC (50:2, protein:enzyme with 0.5 ug/uL enzyme mix) at 37 °C. Peptides were purified with STop And Go Extraction tips^34^.

### Mass spectrometry analysis

The purified peptides were resuspended in buffer A (2% acetonitrile, 0.1% trifluoroacetic acid, and 0.5% acetic acid) and separated on a 75-μm by 50-cm PepMap RSLC EASY-Spray column filled with 2-μm C_18_ reverse-phase silica beads (Thermo Fisher Scientific). Peptides were separated along a linear gradient of 3% to 20% buffer B (80% acetonitrile, 0.5% acetic acid) over a 3-h gradient, followed by a wash with 100% buffer B with a 250-nL/min flow rate using an Easy-nLC 1200 high-performance liquid chromatography system (Thermo Fisher Scientific). Peptides were analyzed on an Orbitrap Exploris 240 hybrid quadrupole-orbitrap mass spectrometer (Thermo Fisher Scientific) operated in data-dependent mode, switching between one full scan and MS/MS scans of abundant peaks with full scans (*m*/*z* 400 to 2,000) acquired with a resolution of 120,000 at 200 *m*/*z*.

### Mass spectrometry data processing

The mass spectrometer output files were analyzed using MaxQuant (version 2.2.2)^12^. The peak list was searched using Andromeda against *T. aestivum* (103,673 sequences, Sept. 15, 2022) and *Gibberella zeae* (14.146, Sept. 15, 2022) from UniProt^14,16^. The following parameters were set: trypsin enzyme specificity with a maximum of two missed cleavages, carbamidomethylation of cysteines (fixed modification), and oxidation of methionine and N-acetylation of proteins (as variable modifications). Spectral matching of the peptides was performed with a false discovery rate (FDR) of 1% identified proteins with a minimum of two peptides for protein identification. The mass tolerance for precursor ions was set to 4.5 ppm and mass tolerance for fragment ions was set to 20 ppm.

### Bioinformatics

Data analysis and visualization were performed with Perseus (version 1.6.2.2) and ProteoPlotter^18,35^. Data was filtered to remove contaminants, reverse peptides, and peptides only identified by site, and intensities were log_2_ transformed. Valid value filtering with proteins identified in at least 3 of 4 replicates in at least one group (general analysis) or within each group (core proteome analysis) was performed followed by imputation with a downshift of 1.8 and a width of 0.3 standard deviations. The data was visualized using principal component analysis (PCA) and proteins with significant changes in abundance between the respective comparisons were identified using volcano plots (Student’s *t*-test, p-value < 0.05; FDR = 5%, S_0_ = 1). Dynamic range plots were generated with protein intensities for each sample and 1D annotation enrichment heat maps defined changes in abundance across defined protein categories by Student’s *t*-test, p-value < 0.05; FDR = 5%, and score <-0.5, >0.5 ^36^. Proteins were sorted by Gene Ontology (GO) Biological Processes (GOBP) and GO Molecular Function (GOMF), as well as UniProt Keywords. Box plots were generated with GraphPad Prism v9 and a Student’s t-test p-value < 0.05 was used for statistical testing relative to the untreated control. STRING network interaction mapping was used^37^.

## Results

### Experimental design enables fungal chemotype and strain analyses at the protein level

To assess the differences in wheat response to *F. graminearum* strains producing 15ADON or the combination of 15ADON/3ANX with the goal of defining new host defense responses to emerging 3ANX toxins, we established an artificial FHB infection within the field. Specifically, 10 strains of 15ADON and 15ADON/3ANX chemotypes of *F. graminearum* were cultured and prepared as spray inoculum (either as mycelium or spores; Table 1) for *T. aestivum* grown within the field (Fig. 1A). Following three-weeks of field growth, proteomics was performed on the samples and a total of 7,152 proteins were identified with 3,892 wheat and 1,380 fungal proteins detected after valid value filtering (i.e., protein identified in 3 of 4 replicates) (Fig. 1B).

**Figure 1:**
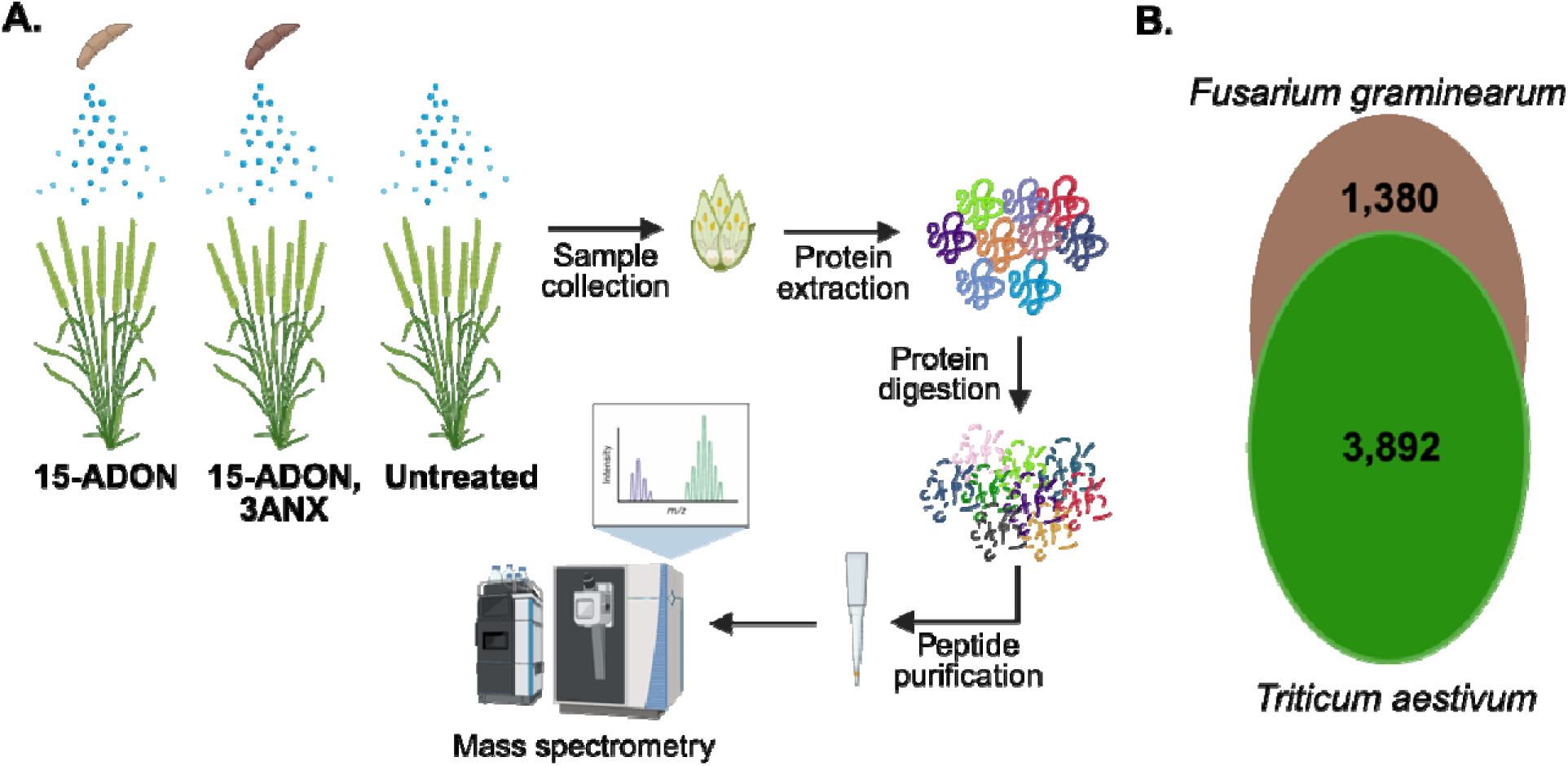
Experimental workflow and proteome overview. **A.** Spray inoculation of *F*. *graminearum* macroconidia from 15-ADON and 15-ADON/3ANX chemotypes was applied on wheat in the field, along with an untreated control. Protein extraction, digestion, and purification performed prior to mass spectrometry-based proteomics profiling. **B.** Number of proteins identified for each biological system after valid value filtering (proteins identified in 3 of 4 biological replicates): *T. aestivum* (green) and *F. graminearum* (brown). Replicates: 10 wheat heads were pooled together from each biological replicate of *F. graminearum* inoculum; four biological replicates of *F. graminearum* were used for inoculum preparation.

### Remodeling of the common wheat proteome is driven by chemotype-specific responses that support defense towards fungal infection

Based on the emergence of *F. graminearum* strains producing 3ANX with increased prevalence and virulence compared to DON^25,29^, we aimed to assess differences in wheat response dependent upon mycotoxin chemotype. For an initial analysis, we focused on the core wheat proteome of 1,398 proteins identified in >70% of all biological replicates within each sample set (i.e., 15ADON/3ANX, 15ADON, control). A principal component analysis (PCA) visualized separation of the samples by treatment (component 1, 21.3%) and biological replicate (component 2, 18.7%) (Fig. 2A). Column correlation values by Pearson correlation to assess replicate reproducibility were 76.3% for control samples, 64.1% for 15-ADON treated samples, and 59.2% for 15ADON/3ANX treated samples (Fig. S1). Assessment of dynamic range of protein intensity spanned >7 orders of magnitude for control samples, >6 orders of magnitude for 15ADON samples, and >5 orders of magnitude for 15ADON/3ANX samples (Fig. 2B). These differences in dynamic range suggest altered protein production (e.g., translation) upon infection with reduced detection of low abundant proteins upon infection with 15ADON/3ANX strains.

**Figure 2:**
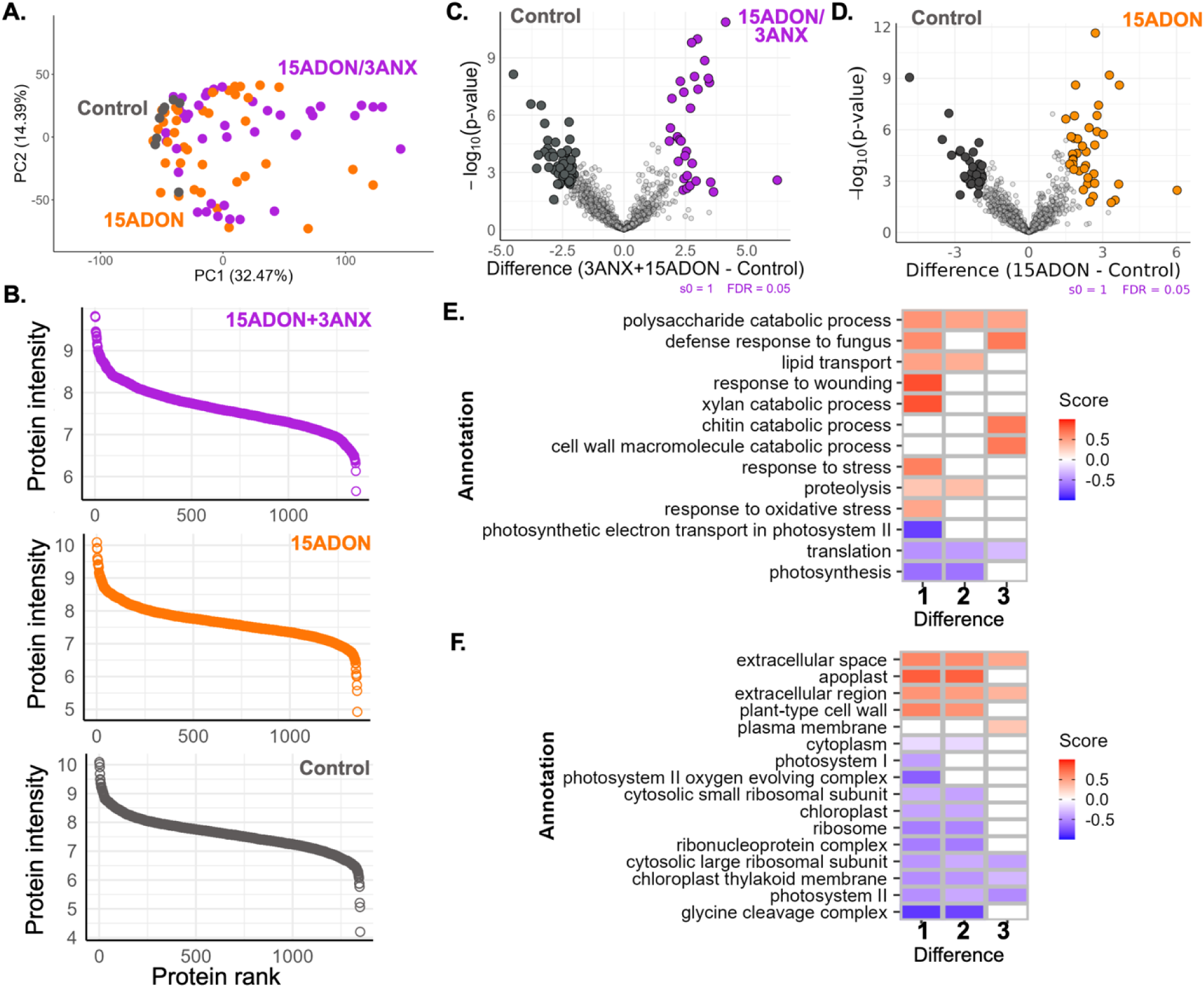
Remodeling of the wheat proteome by mycotoxin chemotype. **A.** Principal component analysis. **B.** Dynamic range of protein intensity by protein rank for each condition. **C.** Volcano plot for differences in protein abundance between 15ADON/3ANX compared to control samples. **D.** Volcano plot for differences in protein abundance between 15ADON compared to control samples. **E.** 1D annotation enrichment by Gene Ontology Biological Processes. **F.** 1D annotation enrichment by Gene Ontology Cellular Compartment. Statistical analysis for volcano plot by Student’s t-test p-value <0.05; FDR = 5%; S_0_ = 1 and 1D annotation enrichment by Student’s t-test p-value <0.05; FDR = 5%; score <-0.5, >0.5. Labels on x-axis for heatmaps: 1 = 15ADON/3ANX vs. control; 2 = 15ADON vs. control; 3 = 15ADON/3ANX vs. 15ADON.

Next, we assessed significant differences in wheat protein abundance driven by mycotoxin chemotype. We observed a significant increase in abundance of 31 proteins compared to a significant decrease in 53 proteins in the 15ADON/3ANX-treated samples compared to controls (Fig. 2C). Similarly, we observed a significant increase in abundance of 38 proteins compared to a significant decrease in abundance of 32 proteins in the 15ADON-treated samples compared to control samples (Fig. 2D). Notably, no significant differences in protein abundance were observed upon comparison of 15ADON/3ANX and 15ADON-treated samples.

Consideration of wheat proteins with significant increases in abundance common to both chemotype-treated conditions identified 29 proteins, including those with known roles in plant defense and stress response (e.g., peroxidases), seed development (e.g., caleosin), environmental sensing (e.g., jacalin-type lectin domain-containing proteins), cell wall thickening (e.g., pectin acyltransferase), and pathogen defense (e.g., ß-amylase) (Table S2)^38–45^. Two proteins were only significantly different upon exposure to the 15ADON/3ANX chemotype, including a fungal lipase-like domain-containing protein and alpha-amylase, with known roles in lipid metabolism and starch degradation and plant defense response, respectively^46,47^. Interestingly, within the 15ADON-treated samples, three leucine-rich repeat proteins, often associated with plant growth, development, and stress response, including immunity against pathogens^48,49^, were detected but absent within the 15ADON/3ANX-treated samples, suggesting possible evasion of immune response detection in the presence of 3ANX.

Lastly, a 1D annotation enrichment heatmap revealed a common enrichment of polysaccharide catabolic processes within chemotype-inoculated samples, whereas an enrichment of translation-associated proteins was observed within the control samples (Fig. 2E). Unique to the 15ADON/3ANX treatment included enrichment of proteins associated with response to wounding, xylan catabolic process, response to stress, proteolysis, and oxidative stress. Further, analysis by GOCC showed common enrichment of proteins associated with the extracellular space and region, as well as enrichment of the apoplast and cell wall upon chemotype treatment, compared to enrichment within the control samples of proteins associated with the cytosol, chloroplast, and photosystem (Fig. 2F). Together, these data demonstrate a general response of wheat to *F. graminearum* infection regardless of mycotoxin chemotype and highlights the activation and subsequent production of proteins in response to the presence of 3ANX.

### Chemotype-exclusive proteins reveal specific host responses to 3ANX

Given our observation of altered wheat protein production profiles dependent upon fungal chemotype treatment, we investigated proteins exclusively detected in the presence of 3ANX. Here, we observed common detection of 1,259 proteins upon 15ADON/3ANX or 15ADON treatment with 32 proteins exclusive to 15ADON/3ANX treatment and 326 proteins exclusive to 15ADON treatment (Fig. 3A). Notably, the detection of a relatively small group of wheat proteins exclusive to 15ADON/3ANX treatment compared to 15ADON treatment suggests that the plant is less adapted to remodel its proteome and defend against the pathogen with the same magnitude of response in the presence of 3ANX. We focused on the 3ANX- exclusive proteins and observed almost 50% were involved in structural and binding activities and endopeptidases with diverse functions, including ß-1,3-glucanase with roles in defense response by hydrolyzing the ß-1,3-glucans of the fungal cell wall, leading to pathogen weakening and death (Fig. 3B; Table S3)^50,51^. Proteins associated with functions in oxidoreductase, defense, and inhibition account for another 30% of the protein IDs, including seed storage domain-containing proteins, which may activate grain filling and associated mechanisms (e.g., cell wall thickening) as a protective mechanism against the mycotoxin^52–54^. Lastly, proteins associated with transport, transferases, and uncharacterized proteins were identified, including a glutathione transferase and glucanotransferase with defined roles in mycotoxin detoxification^55,56^. Together, these data support unique responses of wheat in the presence of 3ANX, including known responses to mycotoxin contamination and defense response, as well as putative novel protein activation mechanisms.

**Figure 3:**
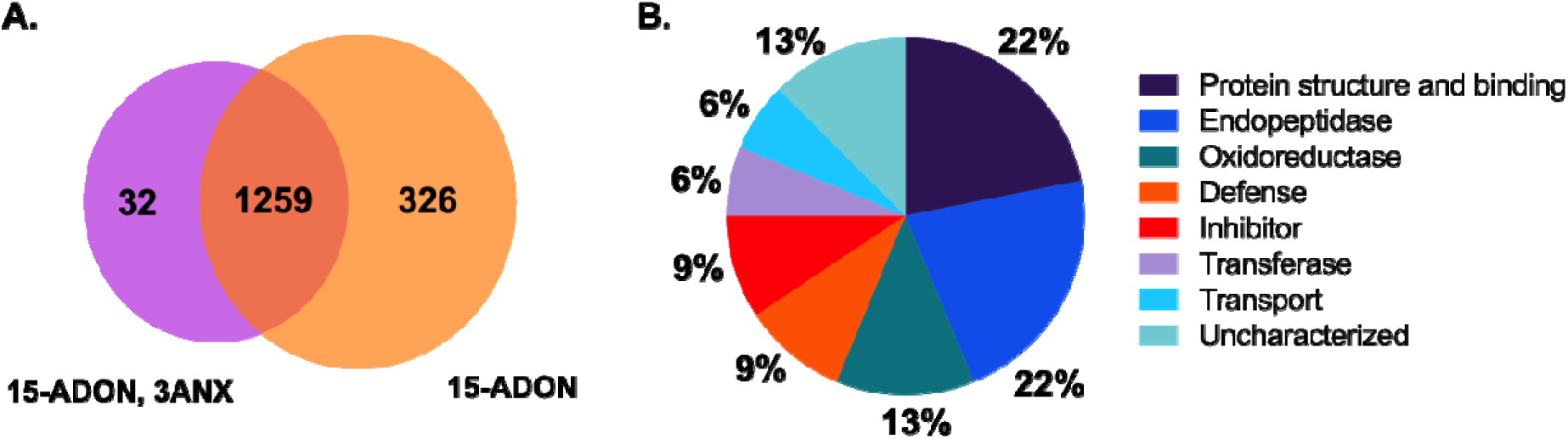
Wheat proteins exclusively detected upon 15ADON/3ANX chemotype treatment. **A.** Venn diagram of proteins common (1,259 proteins) to both chemotype treatments, exclusive to 15-ADON chemotype treatment (326 proteins), and exclusive to 15ADON/3ANX treatment (32 proteins). **B.** Pie chart based on Keywords (UniProt) for proteins exclusive to 15ADON/3ANX.

### Chemotype-specific fungal proteome remodeling emphasizes proteasome and mycotoxin-associated pathway activation

To move beyond the host response and identify putative fungal proteins influenced by the presence and mycotoxin production of the 15DON/3ANX chemotype, we explored the data from the fungal perspective. For instance, a PCA of the common fungal proteome across the chemotypes showed distinction across component 1 (72.36%) and component 2 (6.23%), although, the clusters were not associated with mycotoxin chemotype or known factors of fungal culture source or preparation, suggesting other drivers of fungal proteome remodeling (Fig. 4A). An observation of dynamic range for the fungal proteome highlighted similar detection for both 15ADON/3ANX and 15ADON producing strains covering >4 orders of magnitude in protein abundance (Fig. 4B). We observed 119 fungal proteins exclusively produced by the 15ADON/3ANX chemotype (Fig. 4C; Table S4). A closer look into the functions of these 119 proteins by STRING interaction network mapping and MCL (Markov Cluster Algorithm)-based clustering defined 17 clusters, including an emphasis on proteins associated with the translation, oxidative phosphorylation, metabolism (i.e., phenylalanine sulfur amino acids, acyl-CoA, TCA, cyanoamino acid), as well as the proteasome, mannosyltransferase, and glycosidase hydrolase (Fig. 4D). Importantly, these latter categories have defined roles in cell wall integrity and fungal virulence, suppression of plant immunity, and mycotoxin production^57–59^. A complementary KEGG pathway enrichment plot showcased the involvement of metabolic pathways and the biosynthesis of secondary metabolites, as well as roles of oxidative phosphorylation, glycolysis, and the proteasome (Fig. 4E). Together, these data highlight a chemotype-exclusive proteome in *F. graminearum* in the presence of 3ANX and revealed protein interactions and pathways activated during infection.

**Figure 4:**
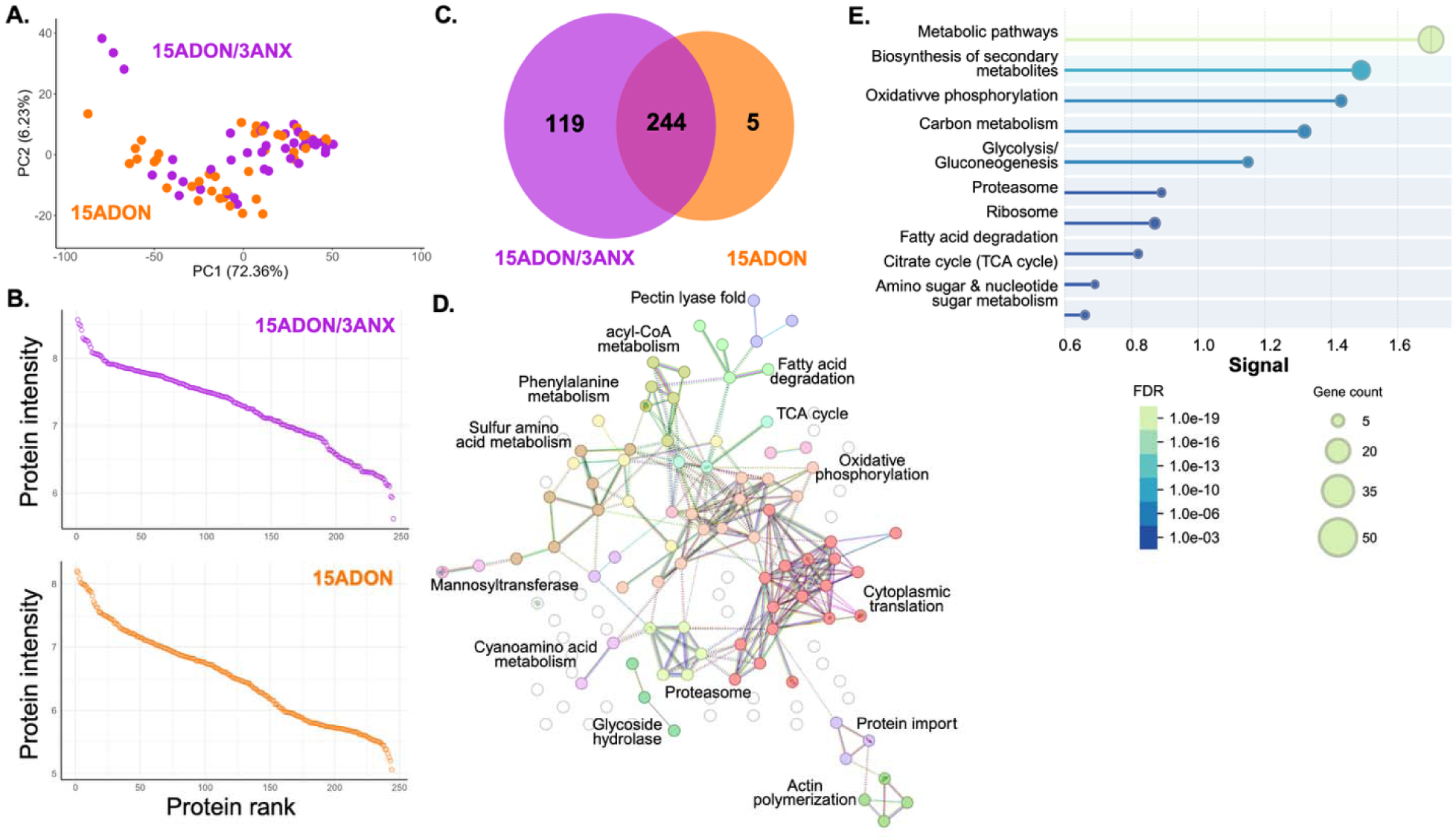
Fungal proteome remodeling by mycotoxin chemotype. **A.** Principal component analysis. **B.** Dynamic range of protein intensity by protein rank for each condition. **C.** Venn diagram of proteins common (244 proteins) to both chemotype treatments, exclusive to 15ADON chemotype treatment (5 proteins) and exclusive to 15ADON/3ANX treatment (119 proteins). **D.** STRING interaction network analysis. MCL clustering (inflation parameter = 3); 17 defined clusters with 14 labeled on image (3 uncharacterized protein clusters not labelled). **E.** KEGG Pathway Enrichment. Terms grouped by similarity >0.8.

### Strain-specific proteome remodeling influences host fungal defenses and mycotoxin protection and alters production of known fungal effectors

Given the differences observed at the protein level across mycotoxin chemotypes, we investigated strain-specific responses from the wheat and pathogen perspectives. For wheat, we identified 686 common proteins across all strains with no proteins unique to any one strain. Among these common proteins, we prioritized proteins with defined roles in pathogenesis (e.g., Pathogenesis-Related proteins), defense response (e.g., chitinases, glucanases), and mycotoxin defenses (e.g., glutathione transferases)^60–62^. Notably, although we noticed differences in abundance of Pathogenesis-Related proteins (i.e., PR, PR1, PR1-1, PR1-2, PR1-19, PR4a, and PR4b), we did not observe a significant increase in production in the presence of the *F. graminearum* strains, indicating that this mechanism of defense was not dominant during the collected stage of infection (i.e., 3 weeks after inoculation) (Fig. S3A). We observed similar trends across the identified chitinases (N=14) with fluctuations in enzyme production but not at a significant level (Fig. S3B). Conversely, a comparison of ß-glucanase production in wheat following inoculation with the *F. graminearum* strains, showed a significant decrease in enzyme abundance in 15ADON-producing strain 12 relative to the untreated control, suggesting a weaker defense response towards this strain (Fig. 5A). Next, a comparison of glutathione transferase production in wheat following inoculation with the *F. graminearum* strains, showed a significant reduction in abundance for two 15ADON/3ANX producing strains (i.e., 13 and 16), indicating a weaker response of the wheat towards these strains, which may be associated with the presence of 3ANX (Fig. 5B).

**Figure 5:**
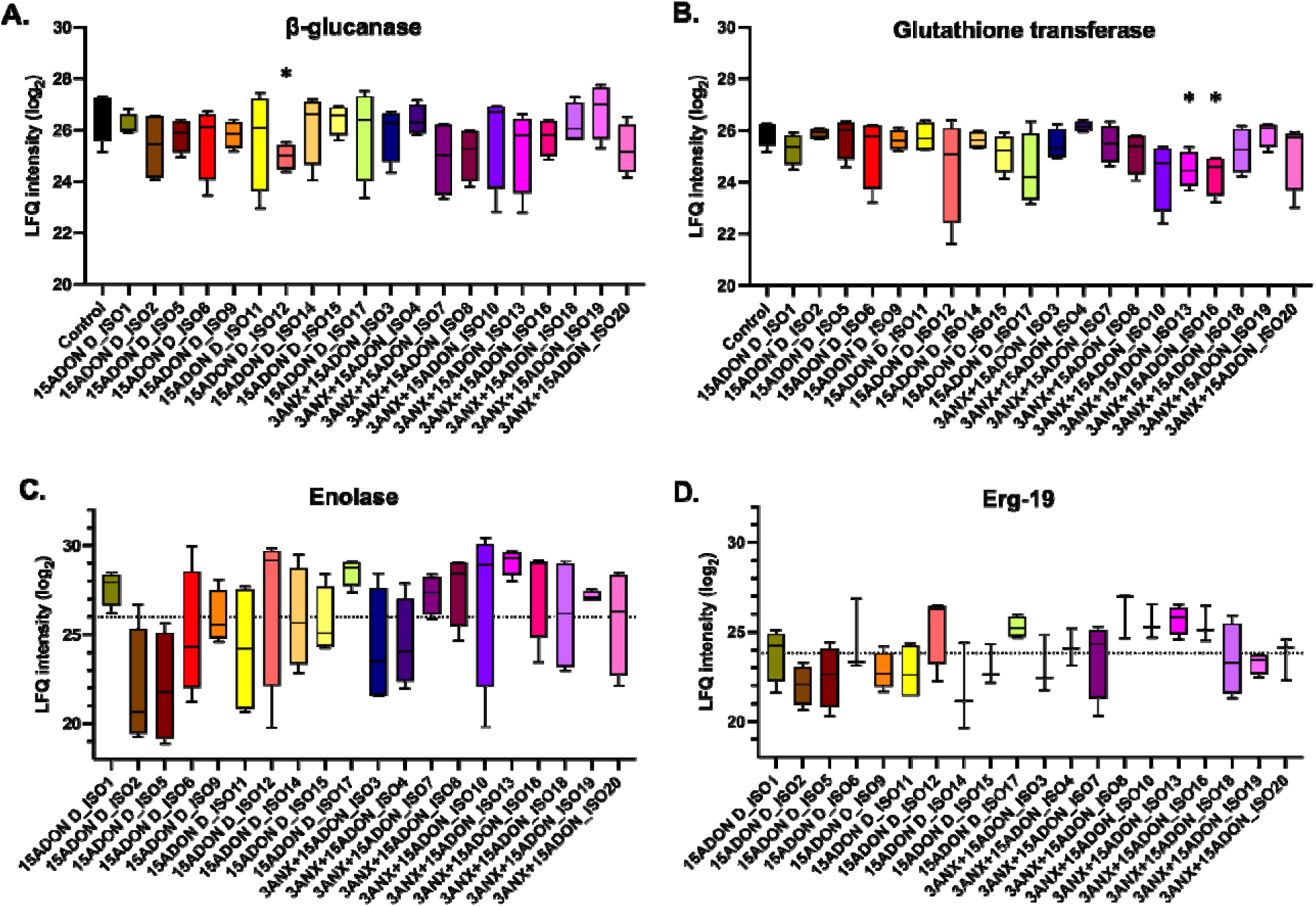
*F. graminearum* strain-specific protein production profiles in wheat and fungi. **A.** Abundance of β-1,3-glucanases from wheat (N = 4). Student’s t-test upon comparison to untreated control, *p-value < 0.05 compared to untreated control (in wheat proteome only). **B.** Abundance of glutathione transferases from wheat (N = 8). Student’s t-test upon comparison to untreated control, *p-value < 0.05 compared to untreated control (in wheat proteome only). **C.** Abundance of enolase from *F. graminearum* (N = 1). Dashed line represents mean of all LFQ intensities across conditions. **D.** Abundance of ergosterol biosynthesis protein 19 from *F. graminearum* (N = 1). Dashed line represents mean of all LFQ intensities across conditions. Mean and standard deviation presented across four biological replicates per condition.

From the fungal perspective, we identified 104 common proteins across all strains and prioritized two proteins: enolase with defined roles in fungal glycolysis contributing to pathogen adhesion and invasion^63^ and an ergosterol biosynthesis protein, Erg19, involved in fungal cell membrane fluidity, permeability, and integrity^64^. Here, we observed fluctuations in protein production across the strains with reduced enolase abundance in 15ADON-producing strains, 2 and 5 (Fig. 5C). For Erg19, one 15ADON-production strain, 17, showed elevated production, whereas six 15ADON/3ANX-producing strains, 4, 8, 10, 13, 16, and 20 showed higher production (Fig. 5D). These data suggest a novel connection between ergosterol biosynthesis and 3ANX production that warrants further investigation to better define the virulence mechanisms used by the fungi during infection.

## Discussion

The recent occurrence of *F. graminearum* strains producing 3ANX and 15ADON/3ANX warrants study into fungal adaptation strategies driving these chemotypes and whether the host crop also has distinct defenses against these novel chemotypes. To tackle this challenge, we applied mass spectrometry-based proteomics of wheat heads collected following field trials inoculated with multiple strains from 15ADON- and 15ADON/3ANX-chemotypes of *F. graminearum*. We also included an untreated control for comparisons. Our study identified specific protein profiles with altered abundance within a core wheat proteome that supported defense response, lipid transport, and cell wall thickening in the presence of the fungi, along with reduced photosynthesis. We also observed the production of specific proteins in samples inoculated with 15ADON/3ANX strains with an emphasis on host metabolism and enzymatic responses, including tissue degradation and reactive oxygen species mitigation. From the fungal perspective, we observed increased abundance of proteins associated with each mycotoxin chemotype, as well as proteins exclusively produced in the presence of 3ANX strains. These proteins included proteasome, mannosyltransferase, and glycosidase hydrolase with defined roles in cell wall integrity and fungal virulence, suppression of plant immunity, and mycotoxin production, likely illustrating the effect of 3ANX on the fungus itself. Finally, by assessing protein abundance profiles for each strain, we observed strain-specific production patterns demonstrating reduced defense response and mycotoxin protection by wheat, and elevated ergosterol production in *F. graminearum* driven by the presence of 3ANX. Overall, this study provides new insight into how the host and pathogen adapt at the protein level to the emergence of the 15ADON/3ANX chemotype and suggests new strategies used to evade plant defense and immune responses, while increasing fungal virulence.

Within our data set, we observed activation of classical wheat defenses, including peroxidases, which are associated with reactive oxygen species modulation and cell wall modifications. Such peroxidases play an essential part in altering the plant’s physical barriers, making it more resistant to pathogen invasion and colonization^64^. We also detected elevated abundance of Caleosin/Peroxygenases *(CLO/PXGs)* as multifunctional enzymes in plant biotic stress responses. These enzymes are involved in detoxification of xenobiotics, synthesis of antifungal compounds, production of cutin monomers integral to plant cuticles for defense against pests and disease, and oxylipin biosynthetic pathways that produce oxidized polyunsaturated fatty acids to mediate plant responses to biotic stress^65,66^. More recently, the possible roles of CLO/PXGs in plant responses to environmental toxins, such as dioxins, have been explored, due to increasingly diverse roles in oxylipin metabolism and other signalling functions^67^. We also defined an increasing abundance of proteins involved in cell wall thickening, such as pectin acyltransferase, to promote strengthening of the cell wall as a major physical barrier against fungal pathogens, including biotrophs, hemibiotrophs, and necrotrophs, to enhance resistance to disease^68^. Taken together, our findings emphasize the importance of both enzymatic and structural defenses in wheat response to different strains of *F. graminearum* producing specific mycotoxin chemotypes. Collectively, this research provides insights into the complex interplay of biochemical pathways and physical barriers that enable wheat plants to combat fungal infections, and underscore the importance of understanding these mechanisms for developing more disease resistant varieties through biotechnological interventions and breeding strategies.

Based on our results, we did not observe any significant differences in PR-protein production across the *F. graminearum*-infected samples compared to the controls. Following *Fusarium* spp. infection in wheat, elevation of PR proteins and gene transcripts increase through early activation of genes encoding peroxidase, PR-1, PR-2 (β-1,3-glucanase), PR-3 (chitinase), PR-4, and PR-5 (thaumatin-like protein) in response^69,71^. Although PR proteins are widely recognized as early responders in plant-pathogen interactions, our observed production patterns after infection has progressed for three weeks suggests a shift in the plant’s defense mechanisms over time. Instead of relying on PR proteins, wheat plants appear to employ alternative strategies, including mycotoxin-associated mechanisms and grain filling, to combat FHB during the later stages of infection. This transition highlights the dynamic nature of plant immunity, where initial responses are rapidly activated to inhibit pathogen establishment, but longer-term defenses evolve to maintain resistance. Understanding these late-stage mechanisms could pave the way for targeted research into sustainable resistance strategies. Identifying the factors responsible for these shifts, such as seed development and cell wall thickening, as well as mycotoxin tolerance and new approaches to detoxification^70^ and other late-stage proteins, may reveal new opportunities for enhancing wheat resilience against FHB.

Exploring how 3ANX influences pathogenesis and survival of *F. graminearum* is another critical aspect of the data sets as the potential to uncover new pathways or mechanisms driving fungal survival may also identify strategies to weaken the pathogen and overcome infection. For instance, we detected an enolase with general lower abundance in 15-ADON chemotype strains compared to the 15ADON/3ANX suggesting a role for this enzyme in energy metabolism for fungal growth and development. We hypothesize that emerging strains with the 15ADON/3ANX chemotypes may use current strategies used by the fungus to invade the host to better adapt to a changing environment. For example, with a changing climate that influences temperature, humidity, rainfall, etc. along with the wheat host that must adapt, the fungus also needs to alter its survival strategies to maintain effective adhesion and invasion strategies. We propose that a connection between 3ANX chemotype presence and altered fungal enolase may promote improved fungal invasion. Another important element of fungal pathogenesis and survival is the production of ergosterol, a critical component of the fungal cell wall and target of fungicides (i.e., azoles) in the field. Our observation of elevated Erg9 in strains of the 15ADON/3ANX chemotype, may increase production of ergosterol to protect the fungus from chemical defenses used in the field through elevated membrane integrity and stress adaptation. Furthermore, recent work shows that ergosterol and DON biosynthesis pathways share a common immediate precursor, farnesyl pyrophosphate (FPP), implying that increasing production of Erg9 in 15ADON/3ANX producing *F. graminearum* strain may lead to increased DON and its derivatives^72,73^. The novelty of the association between ergosterol levels and 3ANX production also warrants deeper investigation, as it may reveal novel regulatory mechanisms or metabolic trade-offs in 3ANX-producing strains. These insights not only enhance our understanding of fungal biology but also offer potential strategies to mitigate the impact of fungal pathogens producing 3ANX in agriculture. Ultimately, we aim to use the information gathered from this study to inform selected breeding of wheat varieties with improved resistance towards *F. graminearum* producing 15ADON/3ANX chemotypes as well as uncover and characterize new targets within the pathogen for improved fungicide efficacy.

## Conclusion

This study reveals distinct host and pathogen proteomic adaptations in response to *F. graminearum* strains producing the emerging 15ADON/3ANX chemotype. Wheat exhibited changes in defense, metabolism, and stress-related proteins, while fungal strains showed chemotype- and strain-specific protein production linked to virulence, immune evasion, and mycotoxin biosynthesis. These findings highlight the dynamic molecular interactions shaping host-pathogen responses and provide critical insights into the evolving virulence strategies of *F*. *graminearum*. Next steps include a multi-year assessment of these strains in wheat to explore the stability of such responses, as well as expanding to other hosts (e.g., corn, barley) to define crop-specific modulation. Further, building upon this information, we aim to explore strategies to better prepare and adapt wheat varieties to the presence of the emerging 15ADON/3ANX chemotype for improved food security and safety as pathogens, crops, and humans adapt to a changing climate.

## Acknowledgements

The authors thank Dr. Art W. Schaafsma for his invaluable contributions and vision for improvements to fusarium head blight management and critical assessment of mycotoxin production and contamination. We also thank members of the Geddes-McAlister lab for helpful discussions and constructive comments on the study. The authors thank Samanta Pladwig (MSc) and Victor Limay-Rios for technical assistance, and Dr. Dyanne Brewer, Manager of the University of Guelph Advanced Analysis Centre – Mass Spectrometry Facility for operation of the mass spectrometer.

## Author Contributions

D. H. & J.G.-M. conceptualized and designed the study. S.R., & N.A., performed experiments. S.R., J.A.M. & J.G.-M. analyzed data and generated figures and tables. S.R. & J.G.-M. wrote the first manuscript draft. All authors contributed to manuscript preparation and have read and approved the submitted manuscript.

## Data Availability

The proteomics datasets are publicly available through PRIDE Proteomics Exchange: PXD062991.

For review purposes, please use:

**Project accession:** PXD062991

**Token:** UemwtZmhVcaA

Alternatively:

**Password:** GTLKY0A3lOBj

## Supplemental Files

Table S1: Experimental design and *F. graminearum* strains.

Table S2: Wheat proteins with significantly increased differences in abundance upon chemotype treatment versus control.

Table S3: Wheat proteins exclusively detected upon 15ADON/3ANX chemotype treatment.

Table S4: Fungal proteins exclusively detected upon 15ADON/3ANX chemotype treatment.

Figure S1: Wheat proteome column correlation by hierarchical clustering of Pearson correlation across samples.

Figure S2: Fungal proteome column correlation by hierarchical clustering of Pearson correlation across inoculated samples.

Figure S3: *F. graminearum* strain-specific protein production profiles in wheat.

