## Supplemental for "Mycotoxin-driven proteome remodeling reveals limited activation of *Triticum aestivum* responses to emerging chemotypes integrated with fungal modulation of ergosterols"

**Table S1: Experimental design and *F. graminearum* strains.**

| **Chemotype** | **Inoculum** | **Isolate** | **DAOMC identifier** | **Host*** | **Source** | **Sample ID** |
| --- | --- | --- | --- | --- | --- | --- |
| 15ADON | Mycelium | 2 | 177406 |  | AAFC | 3 |
|  |  |  |  |  |  | 40 |
|  |  |  |  |  |  | 63 |
|  |  |  |  |  |  | 76 |
|  |  | 5 | 178149 |  | AAFC | 24 |
|  |  |  |  |  |  | 6 |
|  |  |  |  |  |  | 65 |
|  |  |  |  |  |  | 86 |
|  |  | 6 | 180376 |  | AAFC | 11 |
|  |  |  |  |  |  | 29 |
|  |  |  |  |  |  | 52 |
|  |  |  |  |  |  | 67 |
|  |  | 9 | 180379 |  | AAFC | 31 |
|  |  |  |  |  |  | 53 |
|  |  |  |  |  |  | 73 |
|  |  |  |  |  |  | 8 |
|  |  | 11 | 180377 |  | AAFC | 13 |
|  |  |  |  |  |  | 36 |
|  |  |  |  |  |  | 64 |
|  |  |  |  |  |  | 82 |
|  |  | 14 | 180378 |  | AAFC | 16 |
|  |  |  |  |  |  | 37 |
|  |  |  |  |  |  | 58 |
|  |  |  |  |  |  | 74 |
|  |  | 15 | 170785 |  | AAFC | 17 |
|  |  |  |  |  |  | 27 |
|  |  |  |  |  |  | 47 |
|  |  |  |  |  |  | 85 |
|  | Spore | 1 |  | Stalk | RT | 2 |
|  |  |  |  |  |  | 42 |
|  |  |  |  |  |  | 61 |
|  |  |  |  |  |  | 70 |
|  |  | 12 | 252442 | Corn | RT | 14 |
|  |  |  |  |  |  | 38 |
|  |  |  |  |  |  | 48 |
|  |  |  |  |  |  | 68 |
|  |  | 17 | 252437 | Wheat | RT | 21 |
|  |  |  |  |  |  | 23 |
|  |  |  |  |  |  | 62 |
| 15ADON,3ANX | Spore | 3 | 252446 | Stalk | RT | 4 |
|  |  |  |  |  |  | 44 |
|  |  |  |  |  |  | 55 |
|  |  |  |  |  |  | 80 |
|  |  | 4 |  | Stalk | RT | 84 |
|  |  |  |  |  |  | 30 |
|  |  |  |  |  |  | 5 |
|  |  |  |  |  |  | 51 |
|  |  | 7 | 252440 | Stalk | RT | 10 |
|  |  |  |  |  |  | 35 |
|  |  |  |  |  |  | 57 |
|  |  |  |  |  |  | 69 |
|  |  | 8 | 252441 | Wheat | RT | 25 |
|  |  |  |  |  |  | 60 |
|  |  |  |  |  |  | 81 |
|  |  |  |  |  |  | 9 |
|  |  | 10 | 252445 | Corn | RT | 43 |
|  |  |  |  |  |  | 54 |
|  |  |  |  |  |  | 7 |
|  |  |  |  |  |  | 88 |
|  |  | 13 | 252438 | Stalk | RT | 15 |
|  |  |  |  |  |  | 41 |
|  |  |  |  |  |  | 66 |
|  |  |  |  |  |  | 75 |
|  |  | 16 | 252439 | Corn | RT | 22 |
|  |  |  |  |  |  | 32 |
|  |  |  |  |  |  | 46 |
|  |  |  |  |  |  | 78 |
|  |  | 18 |  | Stalk | RT | 20 |
|  |  |  |  |  |  | 33 |
|  |  |  |  |  |  | 50 |
|  |  |  |  |  |  | 79 |
|  |  | 19 | 252443 | Stalk | RT | 19 |
|  |  |  |  |  |  | 34 |
|  |  |  |  |  |  | 49 |
|  |  |  |  |  |  | 71 |
|  |  | 20 | 252444 | Wheat | RT | 18 |
|  |  |  |  |  |  | 26 |
|  |  |  |  |  |  | 59 |
|  |  |  |  |  |  | 77 |
| Untreated |  |  |  |  |  | 1 |
|  |  |  |  |  |  | 12 |
|  |  |  |  |  |  | 28 |
|  |  |  |  |  |  | 39 |
|  |  |  |  |  |  | 45 |
|  |  |  |  |  |  | 56 |
|  |  |  |  |  |  | 72 |
|  |  |  |  |  |  | 83 |

DAOMC = Canadian Collection of Fungal Cultures

AAFC = Agriculture and Agri-Food Canada, Ottawa

RT = University of Guelph, Ridgetown campus

*Host = original source of pathogen

**Table S2: Wheat proteins with significantly increased differences in abundance upon chemotype treatment versus control.**

|  |  | **Difference^#^** | |
| --- | --- | --- | --- |
| **Protein ID*** | **Protein name** | **15ADON/ 3ANX** | **15ADON** |
| A9U8G4 | Alcohol dehydrogenase 1 | n/a | 1.66 |
| A0A3B6TUQ0 | Alpha-amylase | 2.21 | n/a |
| A0A3B6IYD4 | Beta-amylase | 2.56 | 2.68 |
| A0A3B6KSH4 | Beta-amylase | 2.68 | 2.61 |
| A0A3B6C9M9 | Caleosin | 2.44 | 2.33 |
| A0A3B6TML5 | Carbonic anhydrase | 3.43 | 3.03 |
| A0A3B6IJ76 | Cupin type-1 domain-containing protein | 6.24 | 6.02 |
| A0A3B6JCJ8 | Cupin type-1 domain-containing protein | 3.53 | 3.65 |
| A0A3B6SQ42 | Desiccation-related protein | 2.76 | 2.70 |
| A0A3B5Y7F7 | FAD-binding PCMH-type domain-containing protein | 3.29 | 2.77 |
| A0A3B6A203 | FAD-binding PCMH-type domain-containing protein | 4.14 | 3.68 |
| A0A3B5XWZ9 | Fungal lipase-like domain-containing protein | 2.05 | n/a |
| A0A3B6HU03 | Gamma-interferon-inducible lysosomal thiol reductase | n/a | 3.51 |
| A0A3B6I054 | Germin-like protein | 3.48 | 3.27 |
| A0A3B6MY89 | GH10 domain-containing protein | 2.93 | 2.49 |
| A0A3B5Y2X5 | GH18 domain-containing protein | 2.18 | 1.95 |
| A0A3B5ZWW3 | GH18 domain-containing protein | 2.29 | 1.90 |
| I6QQ39 | Globulin-3A | 3.64 | 3.35 |
| A0A077RFQ1 | Hypothetical protein | n/a | 1.84 |
| A0A3B5XSW2 | Jacalin-type lectin domain-containing protein | 3.01 | 2.83 |
| A0A3B5YR46 | Jacalin-type lectin domain-containing protein | 2.54 | 2.68 |
| A0A3B5XTW6 | Laccase | 2.87 | 2.77 |
| A0A3B5ZRE3 | Leucine-rich repeat-containing N-terminal plant-type domain-containing protein | 2.48 | 2.31 |
| A0A3B6NLZ1 | Leucine-rich repeat-containing N-terminal plant-type domain-containing protein | n/a | 1.80 |
| A0A3B6PJP3 | Leucine-rich repeat-containing N-terminal plant-type domain-containing protein | n/a | 1.79 |
| A0A3B6SQT9 | Leucine-rich repeat-containing N-terminal plant-type domain-containing protein | n/a | 1.79 |
| A0A3B6ES48 | Pectin acetylesterase | 2.45 | 2.39 |
| A0A3B6HUB8 | Peroxidase | 2.70 | 2.18 |
| A0A3B6QKW7 | Peroxidase | 2.27 | 2.05 |
| A0A3B6TWE0 | Peroxidase | 1.89 | 1.72 |
| A0A3B6QHW1 | Phytocyanin domain-containing protein | 2.99 | 2.45 |
| A0A3B6QKV4 | PLAT domain-containing protein | n/a | 1.50 |
| A0A3B5ZT42 | Serine protease EDA2 | 2.41 | 2.64 |
| A0A3B6ECD1 | Subtilisin-chymotrypsin inhibitor-2A | n/a | 1.78 |
| W5D700 | Subtilisin-chymotrypsin inhibitor-2A | 1.85 | 1.83 |
| A0A3B6DN86 | Tyrosinase copper-binding domain-containing protein | 1.95 | 1.86 |
| A0A3B6N033 | Uncharacterized protein | 2.57 | 2.22 |
| A0A3B6SHL4 | Uncharacterized protein | 2.32 | 2.26 |
| A0A3B6UCI2 | Uncharacterized protein | 2.77 | 2.40 |
| W5G3Y7 | Uncharacterized protein | n/a | 1.98 |

*Protein ID from UniProt

^#^Difference based on normalized LFQ intensities of proteins upon designated chemotype treatment compared to control. Log_2_ transformed data.

n/a refers to proteins not detected within the given condition.

**Table S3: Wheat proteins exclusively detected upon 15ADON/3ANX chemotype treatment.**

| **Protein ID*** | **Protein name** | **Keyword/Function^#^** | **LFQ Intensity (log2)** |
| --- | --- | --- | --- |
| A0A3B6TMZ7 | 1,4-alpha-glucan branching enzyme | Endopeptidase | 24.19 |
| A0A3B6DA73 | 3-deoxy-8-phosphooctulonate synthase | Metabolism | 22.36 |
| A0A3B6AU50 | 4-alpha-glucanotransferase | Transferase | 21.28 |
| A0A3B5XW08 | AIG1-type G domain-containing protein | Protein structure & binding | 25.52 |
| A0A3B6RKD6 | Amino acid transporter transmembrane domain-containing protein | Transport | 22.83 |
| A0A3B5ZPP6 | Aspartic proteinase | Endopeptidase | 20.87 |
| A0A3B6CGA5 | Barwin domain-containing protein | Protein structure & binding | 24.21 |
| A0A077S0N0 | Beta-1,3-glucanase | Endopeptidase | 23.12 |
| C9E1C1 | Defensin | Defense | 24.25 |
| A0A3B6SHW6 | DOMON domain-containing protein | Uncharacterized | 25.05 |
| P16347 | Endogenous alpha-amylase/subtilisin inhibitor | Inhibitor | 23.80 |
| A0A3B6KKT8 | FAD dependent oxidoreductase domain-containing protein | Oxidoreductase | 22.33 |
| A0A3B6RNB6 | FAD-binding PCMH-type domain-containing protein | Oxidoreductase | 22.41 |
| A0A3B6GNX8 | Glutaminyl-tRNA synthetase | Protein structure & binding | 21.82 |
| A0A3B5ZU59 | Glutathione transferase | Transferase | 25.18 |
| A0A3B6JKD5 | HIT domain-containing protein | Endopeptidase | 23.60 |
| A0A3B6QDX9 | Leucine-rich repeat-containing N-terminal plant-type domain-containing protein | Defense | 23.61 |
| A0A3B6ER29 | NAD-dependent epimerase/dehydratase domain-containing protein | Endopeptidase | 23.23 |
| A0A3B6HVU8 | Non-specific lipid-transfer protein | Transport | 25.19 |
| A0A3B6SQG3 | Peptidase A1 domain-containing protein | Endopeptidase | 23.22 |
| A0A3B6RP55 | Peroxidase | Peroxidase | 22.98 |
| A0A3B6JJ78 | Phosphoenolpyruvate carboxykinase | Protein structure & binding | 22.12 |
| A0A3B6HZM8 | Ribosomal protein | Protein structure & binding | 24.74 |
| A0A3B6KPY9 | rRNA N-glycosylase | Defense | 22.55 |
| A0A3B6JK92 | Seed storage helical domain-containing protein | Inhibitor | 24.61 |
| W5D003 | Seed storage helical domain-containing protein | Inhibitor | 23.37 |
| A0A3B5Z3G6 | Uncharacterized protein | Uncharacterized | 20.13 |
| A0A3B6B6L4 | Uncharacterized protein | Protein structure & binding | 23.21 |
| A0A3B6EBF3 | Uncharacterized protein | Uncharacterized | 25.21 |
| A0A3B6GTM0 | Uncharacterized protein | Uncharacterized | 22.55 |
| A0A3B6KRE9 | Uncharacterized protein | Oxidoreductase | 23.86 |
| W5AP46 | Uncharacterized protein | Protein structure & binding | 22.78 |

*Protein ID from UniProt

^#^Keyword and function derived from UniProt terms.

**Table S4: Fungal proteins exclusively detected upon 15ADON+3ANX chemotype treatment.**

| **Protein ID*** | **Protein name** | **Keyword/Function^#^** | **LFQ Intensity (log2)** |
| --- | --- | --- | --- |
| I1RNL0 | Sphingolipid C9-methyltransferase 2 | Lipid biosynthesis | 24.00 |
| Q4I2J8 | Nascent polypeptide-associated complex subunit alpha | Protein transport | 24.60 |
| Q4I963 | Cofilin | Actin-binding | 24.99 |
| Q4IJT5 | Translationally-controlled tumor protein homolog | Protein biosynthesis | 26.06 |
| V6RG22 | Farnesyl pyrophosphate synthase ERG20 | Lipid biosynthesis | 23.23 |
| A0A098DB19 | Chromosome 1 | Ribosomal protein | 26.74 |
| A0A098DGH9 | Chromosome 2 | N/A | 21.72 |
| A0A098DJU9 | Catalase-peroxidase | Oxidoreductase | 24.35 |
| A0A098DMQ4 | Adenosine kinase | ATP-binding | 27.24 |
| A0A098E4T8 | Phosphoglycerate mutase | Glycolysis | 25.29 |
| A0A0E0RRU5 | Chromosome 1 | Flavoprotein | 23.98 |
| A0A0E0RRY2 | Peptide hydrolase | Aminopeptidase | 25.54 |
| A0A0E0SQB2 | Alpha-L-arabinofuranosidase | Carbohydrate metabolism | 24.40 |
| A0A1C3YHQ4 | Chromosome 1 | Signal | 23.49 |
| A0A1C3YIA8 | Protein phosphatase PP2A regulatory subunit B | WD repeat | 23.19 |
| A0A1C3YIX7 | Chromosome 1 | Metalloprotease | 24.25 |
| A0A1C3YJ05 | Chromosome 1 | Ion transport | 26.44 |
| A0A1C3YJ46 | Leukotriene A(4) hydrolase | Metalloprotease | 24.92 |
| A0A1C3YJX2 | Chromosome 3 | Oxidoreductase | 23.89 |
| A0A1C3YKF1 | Chromosome 3 | RNA-binding | 25.89 |
| A0A1C3YLJ4 | Chromosome 4 | RNA-binding | 24.65 |
| A0A1C3YLZ2 | Chromosome 4 | Oxidoreductase | 24.33 |
| A0A1C3YM35 | Chromosome 2 | N/A | 23.91 |
| A0A1C3YM60 | ATP synthase subunit d, mitochondrial | ATP synthesis | 25.15 |
| A0A1C3YMP0 | Peptide hydrolase | Aminopeptidase | 24.81 |
| I1R9Y4 | ATP synthase subunit 5 | ATP synthesis | 25.34 |
| I1RA28 | Chromosome 1 | Amino-acid biosynthesis | 25.12 |
| I1RA49 | ATP synthase subunit 4 | Ion transport | 25.72 |
| I1RA72 | Tubulin alpha chain | Metal-binding | 24.99 |
| I1RA94 | S-adenosylmethionine synthase | Metal-binding | 24.20 |
| I1RAV5 | Cytochrome b-c1 complex subunit 2, mitochondrial | Electron transport | 25.21 |
| I1RBE6 | Mitochondrial processing peptidase | Metalloprotease | 24.86 |
| I1RBH7 | Chromosome 1 | Coiled coil | 24.25 |
| I1RBK1 | NADH-cytochrome b5 reductase | Flavoprotein | 24.24 |
| I1RBS0 | Chromosome 1 | Protein biosynthesis | 26.99 |
| I1RBS7 | Dipeptidyl peptidase 3 | Metalloprotease | 25.75 |
| I1RC18 | Prohibitin | Membrane | 23.20 |
| I1RC39 | Multifunctional fusion protein | Oxidoreductase | 24.54 |
| I1RC61 | 6,7-dimethyl-8-ribityllumazine synthase | Transferase | 24.14 |
| I1RC95 | Chromosome 1 | ATP-binding | 27.08 |
| I1RCF3 | Chromosome 1 | N/A | 27.28 |
| I1RCJ7 | Eukaryotic translation initiation factor 3 subunit D | Protein biosynthesis | 24.21 |
| I1RCK9 | V-type proton ATPase subunit C | Ion transport | 22.64 |
| I1RCT7 | Chromosome 1 | Pyridoxal phosphate | 23.60 |
| I1RCU8 | Chromosome 1 | Kinase | 23.47 |
| I1RDV8 | Methylmalonate-semialdehyde dehydrogenase | NAD | 23.76 |
| I1RE92 | Chromosome 1 | Transport | 25.11 |
| I1RED5 | 1,3-beta-glucanosyltransferase | Glycoprotein | 25.43 |
| I1RFJ5 | Chromosome 1 | Membrane | 25.85 |
| I1RFQ7 | Eukaryotic translation initiation factor 3 subunit C | Protein biosynthesis | 22.92 |
| I1RFT2 | Eukaryotic translation initiation factor 3 subunit L | Protein biosynthesis | 22.28 |
| I1RG66 | Dihydroxy-acid dehydratase | 2Fe-2S | 25.88 |
| I1RHL0 | Lactase | Glycosidase | 23.99 |
| I1RHQ4 | Chromosome 2 | Protease | 27.27 |
| I1RI43 | Chromosome 2 | Oxidoreductase | 25.33 |
| I1RIM4 | Chromosome 2 | N/A | 27.23 |
| I1RJ06 | Chromosome 2 | N/A | 25.72 |
| I1RJ76 | Chromosome 2 | Hydrolase | 21.80 |
| I1RJL9 | Dolichol-phosphate mannosyltransferase subunit 1 | Glycosyltransferase | 24.18 |
| I1RJP5 | Chromosome 2 | Signal | 27.39 |
| I1RJS2 | Chromosome 2 | WD repeat | 23.97 |
| I1RJS5 | Chromosome 2 | N/A | 26.00 |
| I1RJY7 | Acetyltransferase component of pyruvate dehydrogenase complex | Acyltransferase | 24.34 |
| I1RK55 | Acetyl-CoA C-acyltransferase | Acyltransferase | 24.36 |
| I1RKP1 | Superoxide dismutase | Oxidoreductase | 25.16 |
| I1RKU8 | Chromosome 2 | Coiled coil | 25.88 |
| I1RLY2 | Probable beta-glucosidase G | Glycosidase | 23.43 |
| I1RME3 | Chromosome 3 | Carbohydrate metabolism | 25.23 |
| I1RN08 | Proteasome subunit alpha type | Proteasome | 24.20 |
| I1RN16 | Chromosome 3 | Flavoprotein | 23.09 |
| I1RN88 | Pyruvate dehydrogenase E1 component subunit alpha | Oxidoreductase | 24.03 |
| I1RNM7 | Succinate dehydrogenase | 2Fe-2S | 24.74 |
| I1RQ26 | Proteasome subunit alpha type | Proteasome | 24.22 |
| I1RQC9 | Chromosome 3 | Ribosomal protein | 25.81 |
| I1RQI4 | Chromosome 3 | Lyase | 24.01 |
| I1RQW2 | Non-reducing end alpha-L-arabinofuranosidase | Hydrolase | 23.48 |
| I1RRL5 | Chromosome 4 | N/A | 24.45 |
| I1RRX5 | ATP synthase subunit gamma | ATP synthesis | 25.54 |
| I1RRZ1 | Carboxypeptidase | Carboxypeptidase | 24.57 |
| I1RSJ2 | Phosphomannomutase | Metal-binding | 23.38 |
| I1RSL8 | Importin subunit alpha | Protein transport | 23.24 |
| I1RSQ3 | Chromosome 4 | Ribosomal protein | 26.41 |
| I1RSY7 | Chromosome 4 | Glycosidase | 25.46 |
| I1RUD3 | Chromosome 4 | N/A | 23.74 |
| I1RUR8 | Dihydrolipoyllysine-residue succinyltransferase | Acyltransferase | 23.90 |
| I1RVR7 | Homogentisate 1,2-dioxygenase | Oxidoreductase | 23.33 |
| I1RW24 | Nuclear transport factor 2 | Protein transport | 25.69 |
| I1RW82 | Chromosome 2 | Ligase | 23.82 |
| I1RWK3 | Chromosome 2 | Ribonucleoprotein | 26.18 |
| I1RWM4 | Malate synthase | Peroxisome | 22.48 |
| I1RWP4 | Probable endonuclease LCL3 | Cytoplasm | 23.34 |
| I1RX10 | Chromosome 2 | Lyase | 25.70 |
| I1RXC1 | P-type Na(+) transporter | ATP-binding | 23.81 |
| I1RY27 | Chromosome 4 | Lyase | 25.49 |
| I1RZA7 | Aspartate aminotransferase | Aminotransferase | 23.55 |
| I1RZH9 | Chromosome 4 | Signal | 22.31 |
| I1RZV1 | Chromosome 1 | WD repeat | 23.70 |
| I1RZW8 | Chromosome 1 | Ribosomal protein | 26.53 |
| I1S097 | Chitin synthase 3B | Cell membrane | 22.49 |
| I1S0H2 | S-(hydroxymethyl)glutathione dehydrogenase | Oxidoreductase | 24.91 |
| I1S0M2 | Chromosome 1 | Proteasome | 23.73 |
| I1S0X6 | Chromosome 1 | Ribosomal protein | 26.34 |
| I1S1W4 | Chromosome 3 | Proteasome | 25.64 |
| I1S1Z9 | Chromosome 3 | N/A | 24.56 |
| I1S202 | Chromosome 3 | Aspartyl protease | 25.85 |
| I1S210 | Chromosome 3 | Flavoprotein | 23.66 |
| I1S269 | Alcohol dehydrogenase | Metal-binding | 25.83 |
| I1S270 | Actin-related protein 3 | Actin-binding | 24.14 |
| I1S2N6 | Feruloyl esterase C | Carbohydrate metabolism | 25.63 |
| I1S3S2 | Chromosome 3 | Carbohydrate metabolism | 22.41 |
| I1S444 | Chromosome 1 | Signal | 25.75 |
| I1S464 | NADH dehydrogenase 1 beta subcomplex subunit 9 | Acetylation | 24.37 |
| I1SAK0 | Chromosome 3 | Glycosidase | 23.89 |
| V6QW22 | 60S ribosomal protein L36 | Ribosomal protein | 27.13 |
| V6R1K4 | glutamate--tRNA ligase | Aminoacyl-tRNA synthetase | 21.92 |
| V6RAM6 | Chromosome 2 | Oxidoreductase | 24.09 |
| V6RCG5 | Chromosome 3 | Ribosomal protein | 25.05 |
| V6REA4 | Chromosome 4 | Oxidoreductase | 24.60 |
| V6RK96 | Serine hydroxymethyltransferase | Transferase | 23.88 |

*Protein ID from UniProt

^#^Keyword and function derived from UniProt terms.

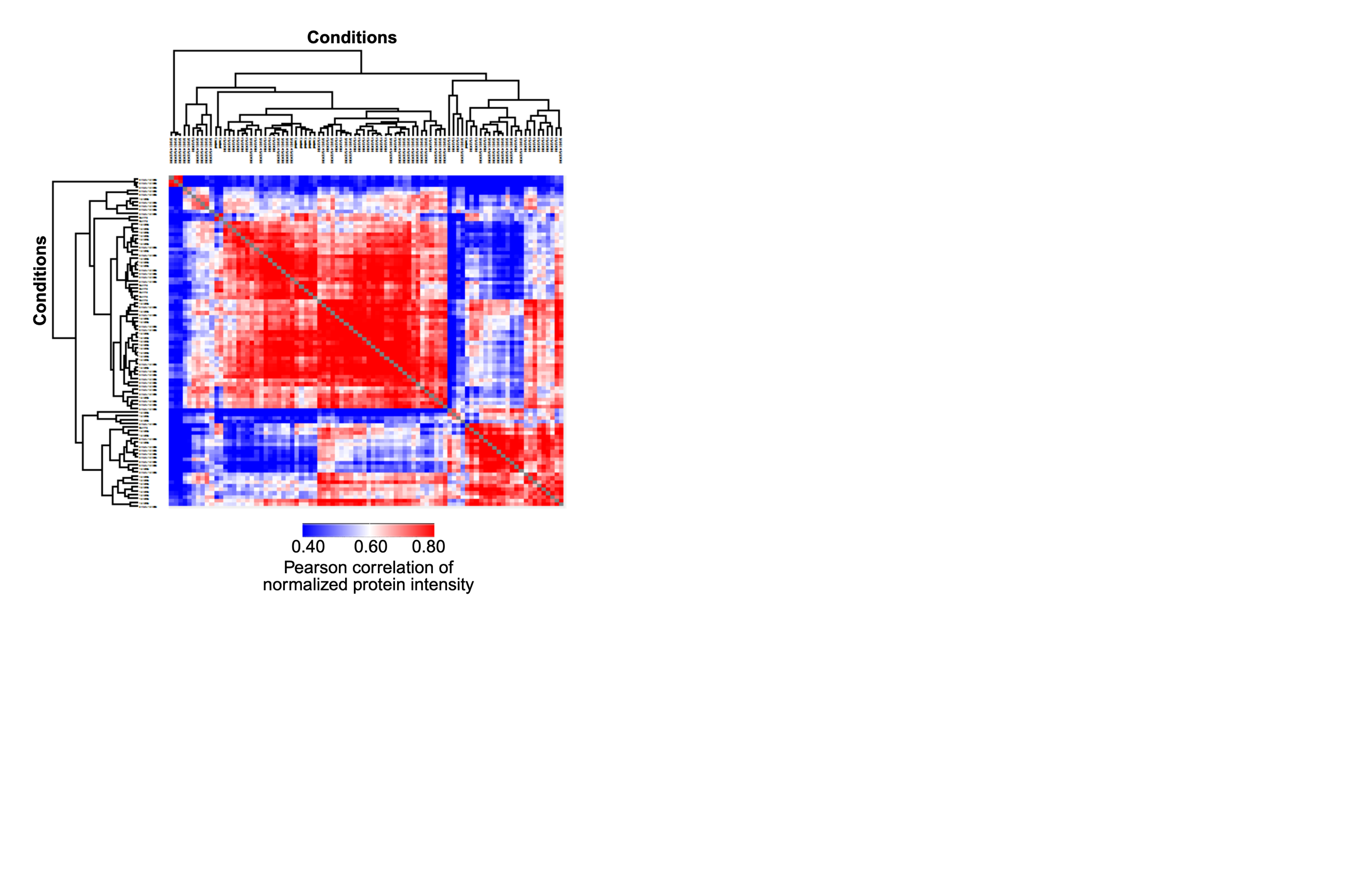

**Figure S1. Wheat proteome column correlation by hierarchical clustering of Pearson correlation across samples.**

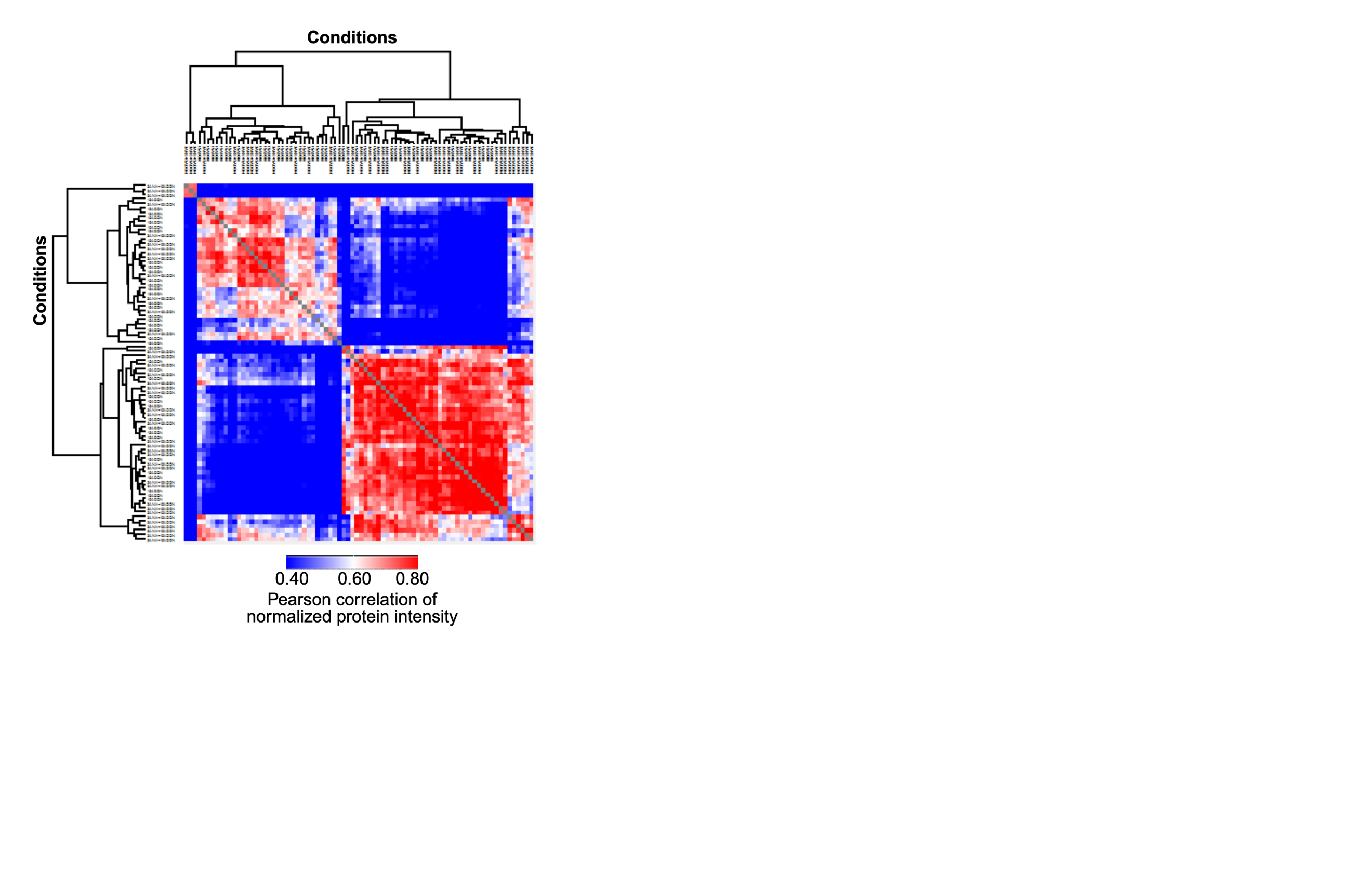

**Figure S2: Fungal proteome column correlation by hierarchical clustering of Pearson correlation across inoculated samples.**

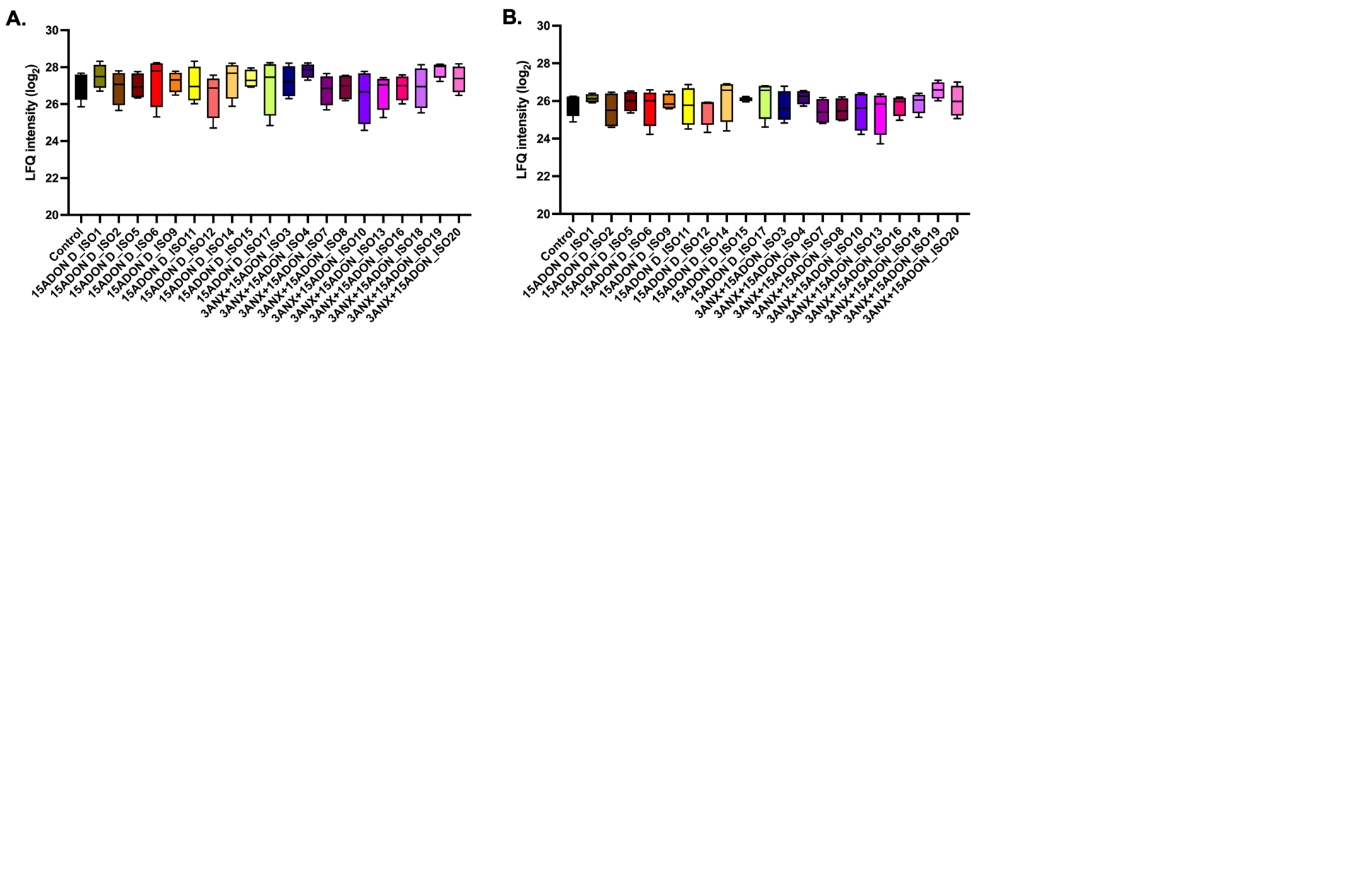

**Figure S3: *F. graminearum* strain-specific protein production profiles in wheat. A.** Abundance of Pathogenesis-related proteins from wheat (N = 7). Student’s t-test upon comparison to untreated control did not define any significant differences. **B.** Abundance of chitinases from wheat (N = 14). Student’s t-test upon comparison to untreated control did not define any significant differences.
